# Choosing an appropriate modelling framework for analysing multispecies co-culture cell biology experiments

**DOI:** 10.1101/008318

**Authors:** Deborah C Markham, Matthew J Simpson, Ruth E Baker

## Abstract

*In vitro* cell biology assays play a crucial role in informing our understanding of the migratory, proliferative and invasive properties of many cell types in different biological contexts. While mono-culture assays involve the study of a population of cells composed of a single cell type, co-culture assays study a population of cells composed of multiple cell types (or subpopulations of cells). Such co-culture assays can provide more realistic insights into many biological processes including tissue repair, tissue regeneration and malignant spreading. Typically, system parameters, such as motility and proliferation rates, are estimated by calibrating a mathematical or computational model to the observed experimental data. However, parameter estimates can be highly sensitive to the choice of model and modelling framework. This observation motivates us to consider the fundamental question of how we can best choose a model to facilitate accurate parameter estimation for a particular assay. In this work we describe three mathematical models of mono-culture and co-culture assays that include different levels of spatial detail. We study various spatial summary statistics to explore if they can be used to distinguish between the suitability of each model over a range of parameter space. Our results for mono-culture experiments are promising, in that we suggest two spatial statistics that can be used to direct model choice. However, co-culture experiments are far more challenging: we show that these same spatial statistics which provide useful insight into mono-culture systems are insufficient for co-culture systems. Therefore, we conclude that great care ought to be exercised when estimating the parameters of co-culture assays.

## 1 Introduction

*In vitro* cell biology assays are an essential element in the study of the migratory, proliferative and invasive properties of different types of cells [Kramer et al. 2013], and they provide insight into various phenomena including malignant spreading [Van Kilsdonk et al. 2010] and wound healing [Xie et al. 2010]. While many types of such *in vitro* assays involve a mono-culture system of a single cell type, many other applications require the analysis of a co-culture system that investigates the migration and proliferation of multiple cell types or subpopulations. For example, wound healing requires the controlled proliferation and migration of both keratinocytes and fibroblasts as well as a number of complex interactions between these two cell types [Wang et al. 2012].

An example of an *in vitro* mono-culture assay is shown in Figure 1(a)–(c). This is a growth-to-confluence assay involving 3T3 fibroblast cells [Todaro et al. 1963]. In this kind of assay, an initially uniform population of cells is placed on a tissue culture plate and monitored in real time as the individual cells within the population move and proliferate to eventually form a confluent monolayer [Simpson et al. 2013]. Analyzing the rate at which the density of cells increases with time allows us to make inferences about the rate at which the cells proliferate [Simpson et al. 2013], which is an essential component of collective cell spreading. Alternatively, an example of an *in vitro* co-culture assay is shown in Figure 1(d)–(f). This is a growth-to-confluence assay involving two cell types: melanocytes and keratinocytes. In this assay, the two cell types are placed on a tissue culture plate and monitored in real time as cells from both subpopulations move and proliferate, eventually leading to a confluent monolayer of cells containing a mixture of both cell types.

**Fig. 1.**
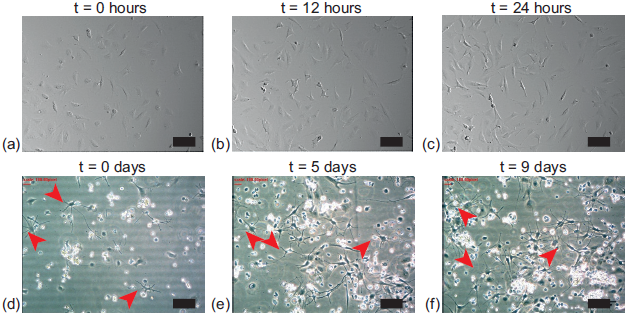
Snapshots showing two growth-to-confluence assays. Images in (a)–(c) show a mono-culture experiment using a population of 3T3 fibroblast cells whereas the images in (d)–(f) show a co-culture experiment containing a subpopulation of melanocyte cells (red arrows) amongst a population of keratinocytes (no arrows). The scale bar in all subfigures corresponds to 100 *µ*m.

Our previous work has focused on developing and applying mathematical models to interpret mono-culture growth-to-confluence experiments [Markham et al. 2013a,Simpson et al. 2013]. In particular, we showed that mono-culture growth-to-confluence experiments can be described using three different mathematical modelling frameworks. Firstly, we considered a stochastic description of individual cell motility, proliferation and death events which has the advantage of directly incorporating individual cell-level behaviours and naturally gives rise to spatial correlations in cell locations, but is analytically intractable [Codling et al. 2008, Deroulers et al. 2009]. Secondly, we considered the traditional corresponding mean-field description of the average cell population density which has the advantage of being analytically tractable but suffers from the disadvantage of neglecting spatial correlations in the distribution of individual cells [Hughes 1995, Liggett 1999]. Thirdly, we considered the more sophisticated corresponding moment dynamics description of the average cell density which has the advantage of being computationally tractable and approximately incorporating the effects of spatial correlations in the distribution of individual cells in the population [Markham et al. 2013b].

Each of these frameworks has been widely employed in the ecology literature to explore the ability of species to invade territories and the potential for disease spread. In particular, the relationship between mean-field approximations and those that incorporate pair-wise interactions have been studied in detail for models of populations undergoing births and death [Dieckmann and Law, Law et al. 2003,Murrell et al. 2004,Raghib et al. 2011], disease spread [Filipe 1999,Filipe and Maule, 2003, Sharkey et al. 2006] and plant dispersal [Bolker and Pacala 1997,Bolker and Pacala 1999]. However, only mean-field models have traditionally been employed to study the evolution of populations of biological cells.

Previous comparisons of these three modelling frameworks in the context of modelling cell biology processes showed that all three produce identical results when the rates of cell proliferation and cell death are sufficiently small relative to the rate of cell motility. Under these conditions the growth-to-confluence process takes place without the population developing any significant spatial correlations. Alternatively, under conditions where the proliferation and death rates are sufficiently large relative to the rate of cell motility, the growth-to-confluence process involves significant spatial correlations and the three models can make very different predictions owing to the extent to which each model includes a description of the effects of spatial correlations [Baker et al. 2010]. The key difficulty identified in our previous work was that each of the three models can always be calibrated to averaged experimental density data to produce an estimate of the underlying rates of proliferation and death [Baker et al. 2010,Simpson et al. 2013]. This can be problematic since the calibration process always leads to a model prediction that matches the observed data, yet if the mean-field or moment-dynamics models are applied under inappropriate circumstances it is possible that the parameter estimates derived from them are meaningless. To address this issue we suggested that some measurement of the degree of spatial correlation ought to be considered to help make an informed decision about when different models ought to be applied [Simpson et al. 2014a, Treloar et al. 2014].

Motivated by the importance of co-culture experiments, such as those shown in Figure 1(d)–(f), the present work seeks to extend and generalize our previous study, which focused exclusively on mono-culture growth-to-confluence experiments, to now investigate appropriate modelling frameworks for analyzing co-culture growth-to-confluence experiments. We achieve this generalization by focusing on situations where we consider co-culture experiments with two cell types, which we refer to as cell type *A* and cell type *B*. This means that we always consider a total population composed of two subpopulations, subpopulation *A* and subpopulation *B,* and we anticipate that the general results and conclusions outlined here will also hold for more general cases involving three or more interacting subpopulations. In Section 2 we outline three mathematical descriptions of a growth-to-confluence experiment with two cell types [Markham et al. 2013b], and we show, in Section 3, that all three descriptions produce similar results when the rates of proliferation and death are sufficiently small whereas the three models produce very different results when the rates of proliferation and death are sufficiently large. These comparisons confirm that a simple model calibration procedure could lead to misleading results and in this light we suggest how additional measurements can be made to provide insight into how the most appropriate modelling framework could be chosen for a particular condition. We conclude with a brief discussion of our results in Section 4.

## 2 Modelling methods

In this work we will use three distinct mathematical models to describe the co-culture experiments: (i) a discrete model that explicitly incorporates individual cell behaviour; (ii) a traditional mean-field model that neglects spatial correlations; and (iii) a moment-dynamics model that approximately incorporates the effects of spatial correlations. Each of these models has been described in detail previously [Markham et al. 2013b, Simpson et al. 2009] and so we only provide a brief description of the key features of each of these models here.

### 2.1 Discrete stochastic description

We consider a population composed of two, possibly distinct, subpopulations on a two-dimensional square lattice, with lattice spacing *∆*, and we invoke an exclusion mechanism whereby each lattice site can be occupied by, at most, a single agent [Simpson et al. 2014b]. Individual agents in each subpopulation undergo unbiased nearest neighbour motility, proliferation and death events. The *i*^th^ species has a movement rate per unit time of 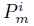 a proliferation rate per unit time of 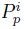 and a death rate of 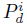 per unit time. To be consistent with a typical experimental scenario, the lattice is initially uniformly populated, at random, meaning that site occupancies are initially uncorrelated and the density is, on average, spatially uniform. When an agent moves or proliferates, the target site is chosen at random from the relevant von Neumann neighbourhood, and the event is aborted if the target site is occupied. We invoke the simplest possible proliferation mechanism whereby agents proliferate to form an identical daughter agent [Simpson et al. 2014b]. Periodic boundary conditions are imposed on all boundaries of the domain. Simulations are propagated in time using a modified form of the Gillespie algorithm [Gillespie 1977] as outlined in [Markham et al. 2013b].

We report results from the stochastic model in two ways. First, we present visual snapshots showing the locations of agents in the population at different points in time. Second, we compute the average agent density across the lattice in the following way: if we are working on an *L*_*x*_ × *L*_*y*_ lattice, then we compute

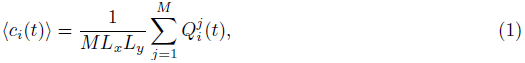

where 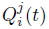 is the number of agents present from subpopulation *i* during the *j*^th^ identically prepared realization of the same stochastic process, and we consider an ensemble of *M* realizations. In this way, 〈*c*_*i*_(*t*)〉 describes the average density of the *i*^th^ subpopulation at time *t.* Since we focus on co-culture experiments with two different cell types, here *i* ∈ [*A, B*]. Sample results from the stochastic model are shown in Figure 2 for a single-species model and in Figure 5 for a two-species model. Throughout this work we take *L*_*x*_ = *L*_*y*_ = 100 and *M* = 100.

### 2.2 Mean-field description

The mean-field description of how the density of each population evolves over time can be written as

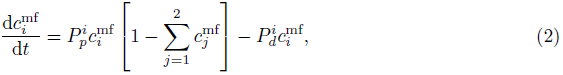

where 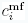 is the density of the *i*^th^ subpopulation and *i* ∈ [*A, B*]. The superscript “mf” explicitly refers to the fact that this model is based on making the traditional mean-field assumption whereby spatial correlations are completely neglected. This model is often referred to as the Lotka-Volterra competition or generalised Verhulst model. The numerical solution of this system of two coupled ordinary differential equations is found using a fourth-order Runge-Kutta method with a constant time step [Chapra et al. 1998].

### 2.3 Moment-dynamics description

The complete details of the derivation of the moment-dynamics description of this system was given previously by us [Markham et al. 2013b] and so here we simply state the main results. The moment-dynamics model describing how the population evolves over time can be written

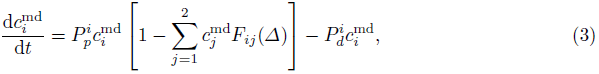

where 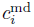 is the density of the *i*^th^ subpopulation and *i* ∈ [*A, B*]. The superscript “md” explicitly refers to the fact that this model is based on an approximate moment-dynamics assumption whereby spatial correlations are approximately incorporated using a moment closure assumption [Markham et al. 2013b]. The *F*_*ij*_(*∆*) term is a correlation function describing the correlation in occupancy of lattice sites at a distance of *∆* between agents of subpopulation *i* and agents of subpopulation *j.* We note that if *F*_*ij*_(*∆*) ≡ 1 then the moment-dynamics description is equivalent to the mean-field description. For a two-species system with the total population being composed of subpopulation *A* and subpopulation *B,* this framework allow us to describe both the autocorrelation for each species, *F*_*AA*_ (*∆*) and *F*_*AB*_ (*∆*), as well as the cross-correlation function, *F*_*AB*_(*∆*), which by symmetry, is equivalent to *F*_*BA*_ (*∆*).

To solve the moment-dynamics model we require equations governing the evolution of *F*_*ij*_(*∆*), and these are generated by considering conservation equations describing the evolution of finding pairs of agents separated by different lattice distances. In turn, these equations require description of the evolution of finding triplets of agents separated by different lattice distances and so on. This means that we have an infinitely large system of conservation equations which we must truncate to obtain an approximate solution. To this end, we close the system at second order using the power-3 Kirkwood superposition approximation (KSA) [Singer 2004]. We choose this closure approximation because, for the models explored here, it performs well over a wide region of parameter space [Baker et al. 2010, Markham et al. 2013b, Simpson and Baker 2011] and, in addition, it ensures positivity. A detailed investigation of the performance of a range of closure approximations was carried out in [Murrell et al. 2004] for an off-lattice, point process logistic-like model. However, a similarly detailed analysis of potential closures for lattice-based exclusion process models such as that explored here lies outside the scope of this work.

On an infinite lattice, the partial differential equations describing evolution of the correlation functions are valid on the domain *∆ ≤ s* < ∞ and are given by

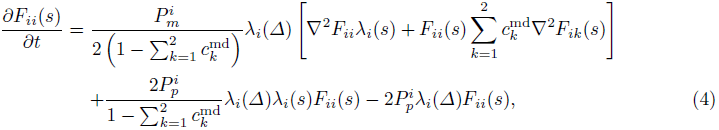

and, for *i ≠ j*,

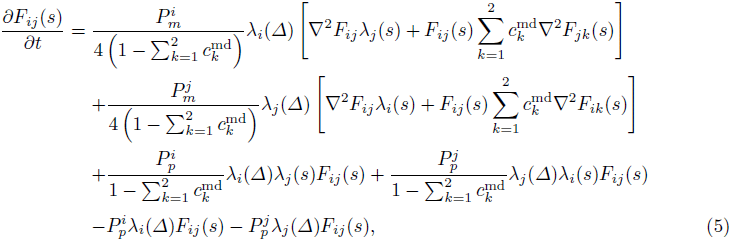

where

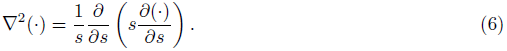

The far-field boundary condition is

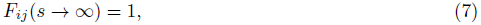

whereas the nearest-neighbour boundary conditions are given by

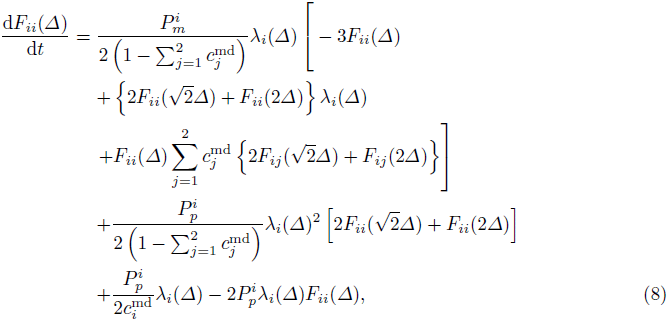

and, for *i ≠ j,*

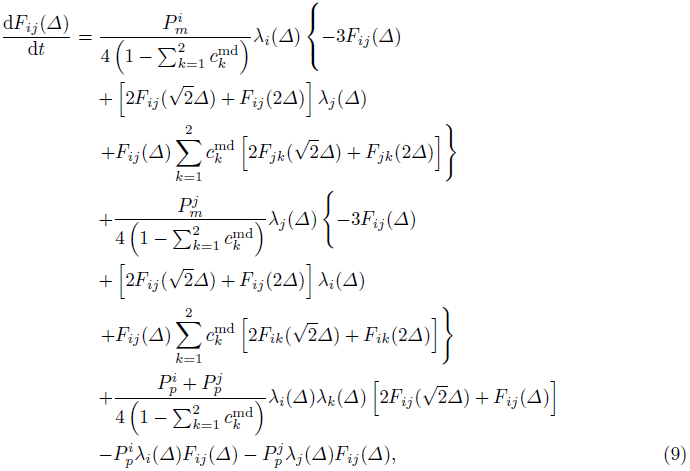

with 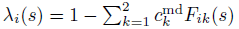.

The numerical solution of this coupled system of nonlinear ordinary and partial differential equations is found by replacing the spatial derivative terms with a central difference approximation on a uniformly discretized domain with mesh size *δs.* The resulting system of coupled nonlinear ordinary differential equations is integrated through time by discretising using the backwards Euler approximation, with time step *δt,* and solving the resulting system of nonlinear algebraic equations using the Thomas algorithm [Press et al. 2007] with Picard linearization and an absolute convergence tolerance of ε [Zheng et al. 2002].

## 3 Results

We will present the results of our study in two sections: first, in Section 3.1, we present results for a single species problem which is relevant when considering a mono-culture growth-to-confluence experiment (Figure 1(a)–(c)). Second, in Section 3.2, we present results for a two species problem which is relevant when considering a co-culture growth-to-confluence experiment (Figure 1(d)–(f)).

### 3.1 Single species results

Results in Figure 2(a)–(c) compare the evolution of the average density profiles predicted by performing repeated stochastic simulations, and averaging the results, compared to the predictions of the mean-field and moment-dynamics models for a single species problem where the initial density of agents on the lattice is 10%. Results in Figure 2(a), corresponding to a high motility rate relative to the proliferation and death rates, show that both the mean-field and moment-dynamics models provide an excellent description of the averaged stochastic data. However, the results in Figure 2(b), corresponding to slightly lower motility rate, show that the mean-field model fails to accurately describe the evolution of the averaged stochastic data, whereas the moment-dynamics model produces results that are visually indistinguishable from the averaged stochastic data at this scale. These results indicate that the additional detail incorporated into the moment-dynamics model lead to an improved prediction under these circumstances. In contrast, the results in Figure 2(c), corresponding to zero motility, indicate that both the mean-field model and the more sophisticated moment-dynamics model can fail to predict the evolution of the average density information. The reason why the mean-field and moment-dynamics models provide different results is because these two model frameworks make different assumptions about the underlying stochastic process. For example, the mean-field description completely neglects spatial correlations whereas the moment-dynamics model approximately neglects spatial correlations, and the accuracy of both descriptions decreases as the rate of proliferation and death increases relative to the rate of migration [Baker et al. 2010, Simpson and Baker 2011].

A key issue that arises when applying a mathematical model to interpret growth-to-confluence experiments, such as the images in Figure 1(a)–(c), is that these images are often converted into a measure of average cell density as a function of time so that data is of the form presented in Figure 2(a)–(c) [Cai et al. 2007,Tremel et al. 2009]. To estimate the proliferation and death rates of the cell population it then seems reasonable to calibrate these parameters using a relevant mathematical model so that the model predictions match the observed data. This kind of calibration approach has been taken by many previous researchers in different contexts including the study of wound healing and malignant spreading [Sengers et al. 2007,Sherratt and Murray 1990, Swanson et al. 2003]. One problem, highlighted by us previously [Baker et al. 2010, Simpson et al. 2013], is that if we take average density information such as that in Figure 2(a)–(c), we can always calibrate either the mean-field or the moment-dynamics model to this data to produce estimates of *P*_*p*_ and *P*_*d*_. However, this commonly-invoked calibration process does not guarantee that the calibrated parameter estimates represent the actual birth and death rates since it is unclear, simply by inspecting density data alone, whether the mean-field and/or the moment-dynamics descriptions are valid. To provide insight into this question of model suitability we need to consider some additional information. For example, the snapshots of the discrete process, shown in Figure 2(d)–(f), illustrate how the agents are arranged on the domain at *t* = 4 for the parameter combinations associated with the density information in Figure 2(a)–(c), respectively. Visual inspection of these images suggests that the agents are arranged relatively uniformly in Figure 2(d), but that there is an increasing degree of spatial clustering and spatial correlation as the birth and death rates increase relative to the motility rate (Figure 2(e)–(f)). To help quantify these differences in spatial clustering and correlation we will now investigate two different quantities.

**Fig. 2.**
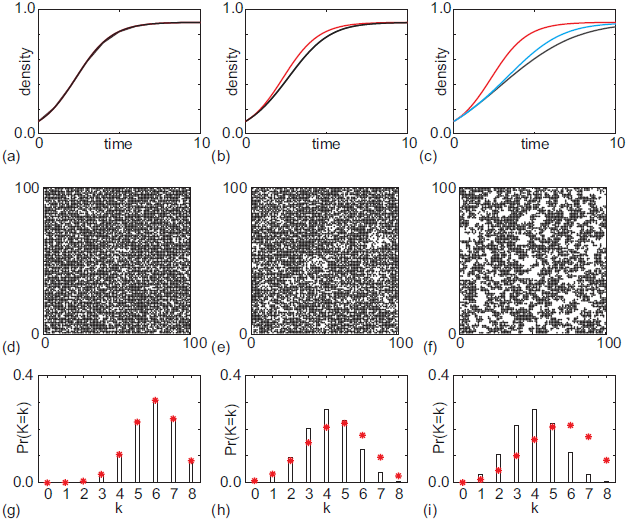
Results for the single species problem as the movement rate, *P_m_*, is varied. *P_m_* decreases from left to right: in the left-hand column *P*_*m*_ = 500; in the centre column *P*_*m*_ = 5; and in the right-hand column *P*_*m*_ = 0. (a)–(c) Comparison of the averaged stochastic results (black) with predictions of the mean-field model (red) and the moment-dynamics model (blue) over time. (d)–(f) Snapshots from a single realisation of the stochastic model at *t* = 4. (g)–(i) Comparison of the averaged agent coordination number (black bars) with the predicted agent coordination number for a spatial uniform system (red asterisks). Parameters are as follows: *P*_*p*_ = 1.0, *P*_*d*_ = 0.1 and initially 10% of lattice sites were occupied.

#### 3.1.1 Agent coordination number

To provide a relatively straightforward measure of the degree of clustering and correlation in the distribution of agents at a particular time we will consider how the agent coordination number distribution, *K,* varies under different conditions. We define the agent coordination number, *K,* for a given site to be the total number of the eight closest sites in the Moore neighbourhood that are occupied, giving *K* ∈ [0, 8]. For a randomly distributed, spatially uniform population at density *C* (*t*) ∈ [0,1], without any clustering or spatial correlations, the average agent coordination number at time *t* will be binomially distributed,

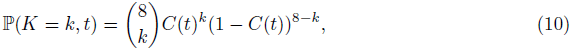

with a mean of 8*C*(*t*) and a variance of 8 *C* (*t*)(1 — C(*t*)) [Simpson et al. 2014a]. This very straightforward expression for the expected distribution of *K* could be compared with experimental or simulation data. In order to obtain an estimate of the agent coordination number from an experimental image, one can impose a lattice on the image, assign each cell to a lattice site, and then count the number of neighbours of each cell.

The results in Figure 2(g)–(i) show, as a series of histograms, the agent coordination number distribution for the systems at *t =* 4 shown in Figure 2(d)–(f). Superimposed on these histograms is the average observed agent coordination number. Comparing the results in Figure 2(g)–(i) we see that: (i) the observed agent coordination number compares well with the binomial distribution, equation (10), when motility is sufficiently high that the mean-field model accurately predicts evolution of the density (Figure 2(a)); (ii) we observe a relatively small discrepancy between the observed agent coordination number and the binomial distribution when the motility is such that the moment-dynamics model captures the observed behaviour but the mean-field model fails (Figure 2(b)); and (iii) we observe a relatively large discrepancy between the observed agent coordination number and the binomial distribution when motility is such that both the moment-dynamics model and the mean-field model fail to capture the observed behaviour (Figure 2(c)).

The qualitative trends indicated in Figure 2 motivate us to consider quantifying these differences. First we define a measure of the difference between the uniform distribution of agent coordination number and the observed distribution of agent coordination number

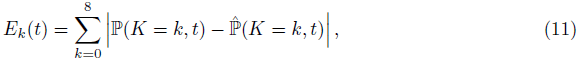

where 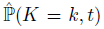 is the averaged observed coordination number distribution at time *t.* To gain an understanding of how the difference in agent coordination number is related to the difference in agent density, we also define

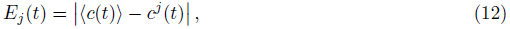

where *j =* mf when we are comparing the performance of the mean-field model, *j =* md when we are comparing the performance of the moment-dynamics model and 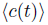 is the averaged observed density at time *t.*

One of the limitations of the results we presented in Figure 2(g)–(i) is that we compare coordination number data at one time point only. For completeness we present additional data, in Figure 3(a)–(b), where we consider single species growth-to-confluence experiments, with the same parameters used in Figure 2(b)–(c), respectively, except now we provide data describing how *E*_k_(*t*) and *E*_*j*_(*t*) evolve with time for both the mean-field and moment-dynamics models. Results in Figure 3(a)–(b) illustrate some key features. Most notably, *E*_k_(*t*) and *E*_*j*_(*t*) vary dramatically with time, and reach some maximum value during the growth-to-confluence process. This means that we ought to be careful as to the particular time we choose to measure *E*_k_(*t*) and *E*_*j*_(*t*), or that we measure both at multiple time points, and we note that the maximum value of *E*_k_(*t*) appears to occur earlier than the maximum value of *E*_*j*_(*t*) in these cases (and all others we investigated).

A summary of results in Figure 3(c) confirms that there is a clear trend between the maximum value of *E*_*k*_(*t*), *E*_kmax_, and the maximum value of *E*_*j*_(*t*), *E*_jmax_, for both the mean-field and the moment-dynamics models. The deviation between the averaged discrete results and both the mean-field and moment-dynamics models increases as *E*_kmax_ increases. Therefore, it is feasible to use a measure of *E*_kmax_ to discriminate between the applicability of the mean-field, moment-dynamics and stochastic descriptions. The discrepancy for the mean-field model increases much more rapidly than the discrepancy for the moment-dynamics model. To illustrate how this kind of information might be used to distinguish between the suitability of the mean-field and moment-dynamics descriptions of this kind of process we have also included dashed horizontal lines in Figure 3(c) to show 1% and 5% thresholds in the discrepancy of the density data. For example, if we were content to use a model that provided no more than a 5% deviation between the averaged stochastic density data and the predictions of the model, then if the observed *E*_kmax_ was less than approximately 0.25 a mean-field model would be suitable, whereas if the observed *E*_kmax_ was between approximately 0.25 and 0.7, a moment-dynamics description would be suitable. If we observed *E*_kmax_ > 0.7, then we could conclude that neither the mean-field or moment-dynamics descriptions were relevant and we ought to focus on using repeated stochastic simulation to interrogate our experimental data.

**Fig. 3.**
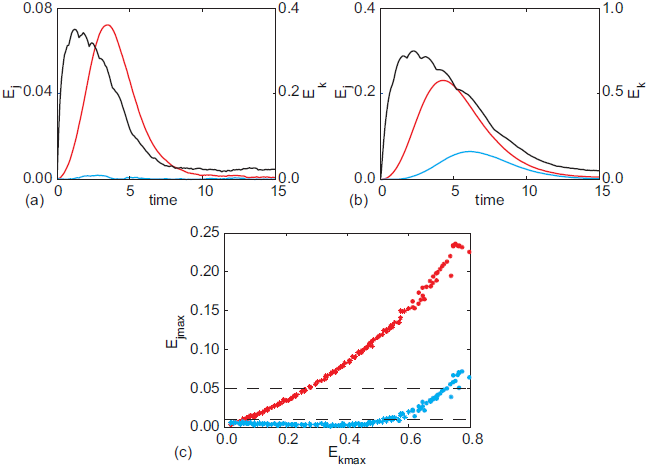
A comparison of the errors between model and data for the single-species system. (a),(b) Evolution of *E*_*j*_(t) for the mean-field model and the averaged stochastic data (*j* = mf, red), and for the moment-dynamics model and the averaged stochastic data (*j* = md, blue) together with evolution of *E*_*k*_(*t*) (black). Parameters are as in the middle and left-hand columns of Figure 2, respectively. (c) The maximum value of *E*_*j*_(*t*), *E*_jmax_, plotted as a function of the maximum value of *E*_k_(*t*), *E*_kmax_, for *j* = mf (red) and *j* = md (blue). The parameter ranges analysed are *P*_*m*_ ∈ [0, 500], *P*_*p*_ ∈ [0, 10], *P*_*d*_ ∈ [0, *P*_*p*_] and the results from simulations using 140 different parameter combinations are displayed. The dashed horizontal lines represent maximum discrepancies (*E*_jmax_) of 0.01 and 0.05 in the density data.

#### 3.1.2 Spatial correlation index

As a second measure of the degree of clustering, we explore how the average correlations in lattice site occupancy for various lattice site distances vary with time. To obtain *F*_*AA*_ (*x*, *t*) we count the number of pairs of agents separated by a distance *x,* and then divide this number by the square of the total number of agents on the lattice [Markham et al. 2013a, Treloar et al. 2014]:

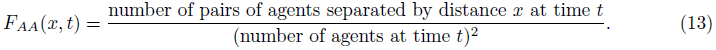

Figure 4(a) compares evolution of the correlation functions at distances *∆,* 2*∆* and 3*∆* as a function of time against evolution of the error, *E*_*j*_(*t*), between both the mean-field and moment-dynamics models and the averaged stochastic data. As with the agent coordination number, there is a delay between the maxima of the correlation function and the maximum error for both models. This indicates that we cannot rely on snapshots of the correlation functions to determine the suitability of a particular model. As for the agent coordination number, we then explored how *E*_jmax_ depends on the maximum of *F*_*AA*_ (*x, t*), *F*_AAmax_, over a wide range of parameter values. Our results suggest that the values of *F*_AAmax_ and *E*_jmax_ are closely related for both models, over distances *∆,* 2*∆* and 3*∆*, with the relationship becoming less well-defined as the distance increases (Figure 4(b)–(d)). The strongest trend corresponds to max{*F* _AA_(2*∆*) - *F* _AA_(*∆*)} and *E*_jmax_ (Figure 4(e)). Therefore, as an alternative to the agent coordination number data, described in Section 3.2.1, we suggest that estimates of max{*F*_AA_(2*∆*) - *F* _AA_(*∆*)} could be used as a measure to distinguish between the suitability of the mean-field and moment-dynamics models in recapitulating the averaged discrete data for mono-culture growth-to-confluence experiments.

### 3.2 Two species results

We now focus on two species systems, which are relevant to co-culture growth-to-confluence experiments, like that shown in Figure 1(d)–(f). Figure 5(a)–(c) shows results corresponding to simulations of a two species system, where the initial density of each species is 5%, so that the total population initially occupies 10% of the lattice, just like in Section 3.1 for the single species problem. Although both species *A* and *B* have the same proliferation rates, the evolution of the two subpopulations is very different with subpopulation *A,* which has a higher movement rate compared to subpopulation *B*, occupying a greater percentage of lattice sites at later times. In Figure 5(a) we plot evolution of the averaged discrete density profiles alongside the results from the mean-field and moment-dynamics models. We see that the mean-field model incorrectly predicts that the two subpopulations will co-exist with each subpopulation eventually occupying 50% of the lattice sites. The moment-dynamics model correctly predicts both the transient and steady state behaviour of the system: the species coexist but subpopulation *A* dominates, occupying around 55% of lattice sites. The snapshots shown in Figure 5(b)–(c) of a single realisation at *t* = 3 and *t* = 10 indicates that the failure of the mean-field model may be due to the emergence of spatial correlations in lattice site occupancy, as was the case for the single-species model, since the build up of clustering in both subpopulations is visually distinct. Further investigation into the failures of the mean-field and moment-dynamics models in various regions of parameter space indicate, as for the single-species model, that the validity of the model assumptions (the independency of lattice site occupancies and the validity of the KSA, respectively) are key to the success of the models in replicating the behaviour of the discrete system [Markham et al. 2013b].

**Fig. 4.**
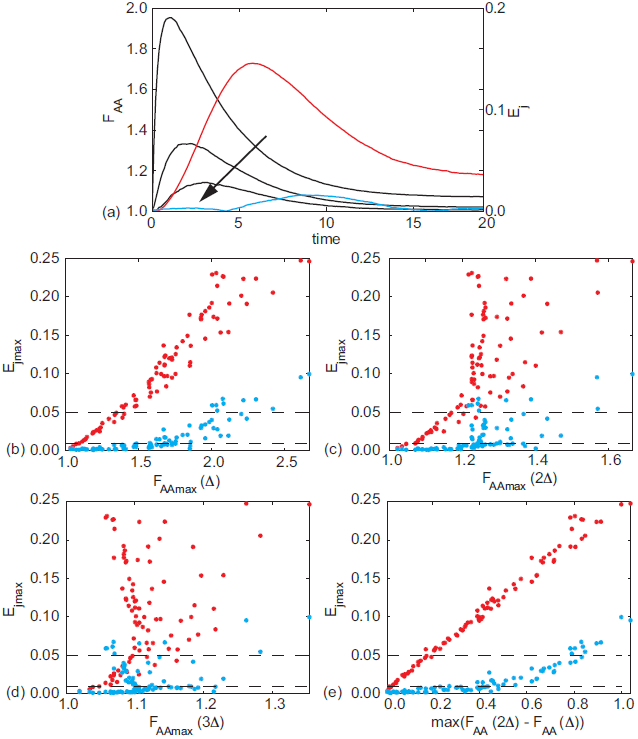
Evolution of the correlation functions. (a) Evolution of the correlation functions at distances *x* = *∆*, *x* = 2*∆* and *x* = 3*∆* as a function of time (black lines), with the direction of increasing *x* indicated by the arrow. These results are superimposed on profiles showing *E*_*j*_(*t*) for the mean-field model (red) and *E*_*j*_(*t*) for the moment-dynamics model (blue). Parameters are as in the middle column of Figure 2. (b)–(e) Plots of *E*_jmax_ as a function of the maximum of *F*_*AA*_ (*t*), *F*_max_, with results from the mean-field model shown in red and from the moment dynamics model in blue. The parameter ranges analysed are *P*_m_ ∈ [0, 500], *P*_*p*_ ∈ [0, 10], *P*_*d*_ ∈ [0, *P*_*p*_] and the results from simulations using 90 different parameter combinations are displayed. As in Figure 3, the dashed horizontal lines represent maximum discrepancies (*E*_jmax_) of 0.01 and 0.05 in the density data.

**Fig. 5.**
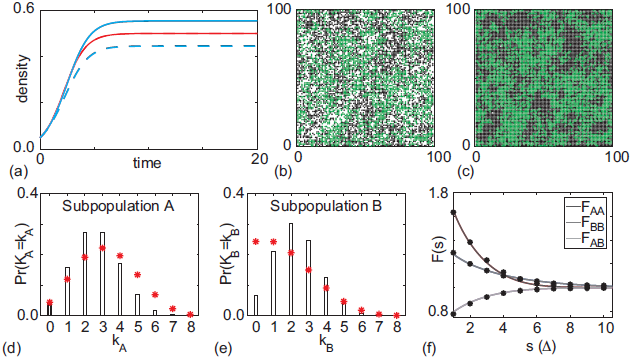
Results for a two species problem. (a) The mean-field model (red) predicts both species evolve to occupy, on average, 50% of lattice sites, whilst the moment-dynamics model (blue) correctly predicts the density of species *A* (solid line) to be greater than that of species *B* (dashed line). In each case the averaged discrete results (black) are obscured by the predictions of the moment-dynamics model. (b),(c) snapshots of the populations at *t* = 3 and *t* = 10, respectively, indicate that each of the species displays significant clustering. (d),(e) Comparison of the averaged agent coordination number (black bars) with the predicted agent coordination number for a spatial uniform system (red asterisks) at *t* = 3. (f) The averaged auto- and cross-correlation functions at *t* = 3. The solid curves correspond to the solution of the moment-dynamics model whereas the dots correspond to averaged simulation data from the stochastic description. Parameters are as follows: 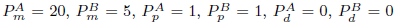 and each species initially occupies 5% of lattice sites uniformly at random.

As with the single species system, it is possible to calibrate either the mean-field or the moment-dynamics descriptions to a given set of averaged stochastic density data to provide an estimate of the model parameters governing the birth and death rates of both subpopulations in this context. However, as before, merely fitting different models may give rise to different parameter estimates, and it is unclear, *a priori*, which modelling framework is the most appropriate when we are dealing only with averaged density information, as is often the case in experimental cell biology [Cai et al. 2007, Tremel et al. 2009]. To provide further guidance, we now investigate whether it is possible to use multi-species equivalents of the agent coordination number and spatial correlation index summary statistics to distinguish between different regions of parameter space in which the mean-field and moment-dynamics models are able to reliably replicate the averaged stochastic data.

### 3.2.1 Agent coordination number

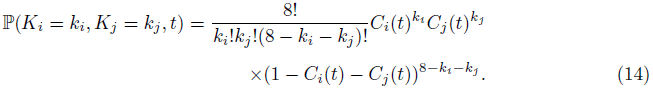

This means that the agent coordination number of each individual subpopulation is binomially distributed,

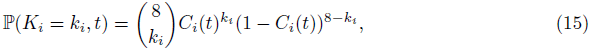

as in the single species case. Results in Figure 5(d)–(e) show the observed averaged agent coordination numbers for the system shown in Figure 5(a)–(c) compared with the theoretical results for the case where the subpopulations are randomly distributed. Note that, had the mean-field model predicted the observed average agent density data in Figure 5(a) correctly, then we would expect to see the actual distribution of agent coordination number match with the expected binomial result, similar to what we saw in Figure 2(g). In contrast, results in Figure 5(d)–(e) indicate that the observed distribution of agent coordination number deviates significantly from the binomial distribution, which is consistent with the fact that the assumptions underlying the mean-field model are not met.

**Fig. 6.**
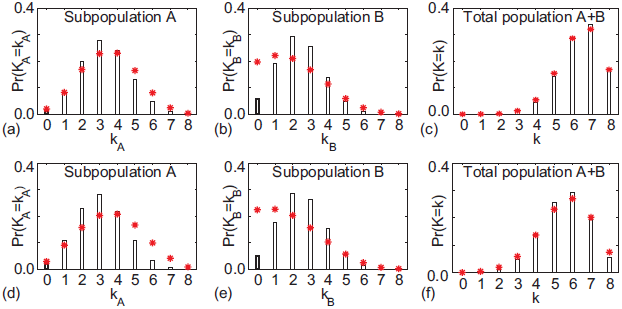
Comparison of the averaged agent coordination number (black bars) with the predicted agent coordination number for a spatial uniform system (red asterisks) for two co-culture experiments. (a)–(c) In isolation, the agent coordination number of species A is close to that theoretically predicted for a uniform population, but this is not the case in co-culture. Parameters are as follows: 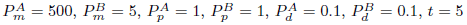 and both species initially occupy 5% of lattice sites uniformly at random. (d)–(f) Each individual species displays an agent coordination number that deviates significantly from that predicted for a uniform population, but this is not the case when they are considered as a single species. Parameters are as follows: 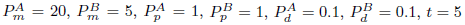 and both species initially occupy 5% of lattice sites uniformly at random.

At first glance, the results in Figure 5 indicate that the agent coordination number may be able to provide a guide as to the suitability of the mean-field and moment-dynamics models in predicting results from the stochastic model in the two-species co-culture model. As a potential means to simplify our calculations, we first ask whether understanding the dynamics of each subpopulation under equivalent mono-culture conditions is sufficient to determine the ability of each modelling framework to replicate averaged stochastic data. Results in Figure 6 compare the observed and predicted distributions in agent coordination number using two different parameter combinations. In Figure 6(a)–(c), subpopulations *A* and *B* have the same parameter values as in the left-hand and centre columns of Figure 2, respectively. This means that, in isolation, we expect the evolution of population *A* to be well approximated by both the mean-field and moment-dynamics models, and that subpopulation *B* will be well-approximated by the moment-dynamics model and poorly approximated by the mean-field model. However, when these two subpopulations are grown together in co-culture, both subpopulations display agent coordination numbers that differ from that expected of a spatially uniform population even though the total agent coordination number indicates that, when considered as a single population, the system does not display significant clustering. This result implies that we cannot draw reliable conclusions regarding the suitability of a particular model for a co-culture experiment simply by considering the isolated behaviour of each of the individual species alone as in a mono-culture experiment.

**Fig. 7.**
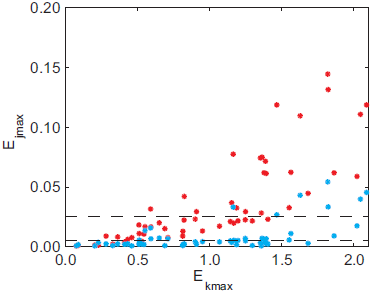
A plot of *E*_jmax_ as a function of *E*_kmax_ over a wide range of parameter space reveals that the agent coordination number is not a suitable metric for distinguishing between the suitability of mean-field (red) and moment-dynamics (blue) models in a co-culture experiment. The parameter ranges analysed are *P_m_* ∈ [0, 1000], *P_p_* ∈ [0,10], *P_d_* = 0 and the results from simulations using 55 different parameter combinations are displayed. As in Figure 3, the dashed horizontal lines represent maximum discrepancies (*E*_jmax_) of 0.01 and 0.05 in the density data.

To provide further insight into modelling multi-species co-culture experiments we now explore whether it is necessary to be able to distinguish between the various subpopulations in order to determine the suitability of one modelling framework over another. Results in Figure 6(d)–(e) demonstrate that there are cases in which the distribution of agent coordination number for each subpopulation differ significantly from that expected from spatially uniform populations. However, in Figure 6(f) we see that if we were unable to distinguish between the subpopulations, and instead we could only determine the total cell density, then calculating the agent coordination number for the total population would erroneously imply that the mean-field modelling framework could be appropriate. Together, these results suggest that, in order to use information about agent coordination number to distinguish between the suitability of the mean-field and moment-dynamics models, it is necessary to measure the agent coordination number of each subpopulation in a co-culture experiment. To deal with this complication, we would aim to produce a plot similar to that shown in Figure 3(c) where we showed that a strong correlation exists between differences in the predicted and observed densities and agent coordination numbers, *E*_jmax_ and *E*_kmax_, respectively, in the co-culture simulations. If this were possible, we could then reliably assume that measurements of *E*_kmax_ for a particular co-culture system would be sufficient to guide our choice of model. Unfortunately, results shown in Figure 7 indicate that the results are more subtle for the co-culture system: we see a much less pronounced relationship between *E*_jmax_ and *E*_kmax_ in the co-culture system compared to the mono-culture system. Therefore, it does not seem reasonable to use the observed average agent coordination number as a means to distinguish the suitability of the mean-field and moment-dynamics models in the context of modelling a co-culture experiment. We now turn to study the spatial correlation index as an alternative summary statistic for co-culture experiments.

### 3.2.2 Spatial correlation index

Finally, we now explore whether the averaged observed correlation functions can improve our ability to make a distinction between the performance of the three modelling frameworks. To estimate *F_ij_*(*x,t*) we count the number of (*i*, *j*) agent pairs separated by a distance x, and normalise by the products of the total numbers of *i* and *j* agents on the lattice [Markham et al. 2013a, Treloar et al. 2014]:

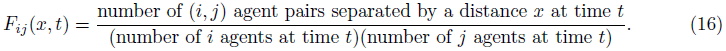

In Figure 5(f) we demonstrate the ability of our moment-dynamics model to approximate both auto- and cross-correlations. As expected, the system shows significant positive local spatial correlations in the distributions of each individual species, and negative local cross-correlations. In Figure 8 we plot *E*_jmax_ as a function of the maxima of the auto- and cross-correlation functions, for a wide range of parameter values. As with the agent coordination number, the results are disappointing in that it does not seem sensible to base our model choice upon this measure of the system correlations in a co-culture experiment.

**Fig. 8.**
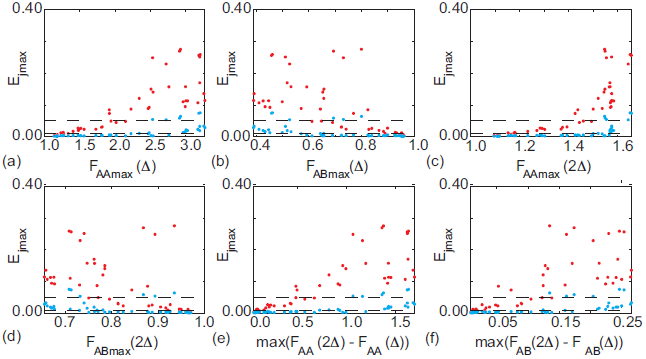
A plot of *E*_jmax_ as a function of the maxima of the auto- and cross-correlation functions over a wide range of parameter space reveals that the agent coordination number is not a suitable metric for distinguishing between the suitability of mean-field (r) and moment-dynamics (blue) models in a co-culture experiment. The parameter ranges analysed are *P*_*m*_ ∈ [0, 1000], *P*_*p*_ ∈ [0, 10], *P*_*d*_ = 0 and the results from simulations using 45 different parameter combinations are displayed. As in Figure 3, the dashed horizontal lines represent maximum discrepancies (*E*_jmax_) of 0.01 and 0.05 in the density data.

## 4 Discussion and conclusion

In this work we explore the performance of three different mathematical modelling frameworks for quantitatively evaluating the results of *in vitro* mono- and co-culture growth-to-confluence assays. In particular, we focus on: (i) a stochastic description of a birth, death, movement process which is analytically intractable but directly incorporates spatial correlation effects, (ii) a traditional mean-field description of a birth-death-movement process which is analytically tractable but completely neglects to incorporate any spatial correlation effects; and (iii) a more sophisticated moment-dynamics description of a birth-death-movement process which is computationally tractable and approximately incorporates the effects of spatial correlations. Since it is relatively common to calibrate such models to experimental data in order to estimate model parameters [Sengers et al. 2007, Sherratt and Murray 1990, Swanson et al. 2003], our aim is to provide a quantitative method which can be employed to distinguish between the validity of each type of model over a wide range of parameter space.

In this work we have used two particular spatial statistical tools, namely agent coordination number and a spatial correlation index, in an attempt to distinguish between the validity of the stochastic, traditional mean-field and moment-dynamics descriptions of a birth-death-movement process. Although we have focused on these two measures, we are aware that there are other types of tests for spatial randomness, such as indices that measure a departure from the complete spatial randomness state [Binder and Landman 2011, Diggle 1983,Phelps and Tucker 2006, Simpson et al. 2013]. Although we did not include any results using this kind of complete spatial randomness index, we did perform a preliminary study using the complete spatial randomness index on the same data presented in this work and found that this method provided no additional insight than the agent coordination number and spatial correlation index method. Furthermore, the complete spatial randomness index is limited in the sense that it depends on partitioning the domain into equally-sized bins and calculating the variance of numbers of objects per bin. Since the results can be quite sensitive to the bin size, and there is little guidance in the literature with regard to selecting the optimal bin size, we found it was computationally expensive to implement this kind of approach since we always had to test whether our results were independent of the bin size.

The results of our study can be summarised in the following way. For mono-culture assays in which we consider just one type of cell, we show that both the averaged agent coordination number and spatial correlation index provide a suitable means of distinguishing between the ability of each model to replicate the observed averaged density data over a wide range of parameter space. While our results for the mono-culture case suggest that we exercise caution against using just one time point during the experiment to measure the agent coordination number or spatial correlation index, we show that either of these spatial summary statistics can be used to reliably distinguish between the application of a mean-field, moment-dynamics or stochastic representation of the growth-to-confluence process when we consider examining the entire time course of the experiment and focus on the maximum deviation between the observed coordination number of spatial correlation index for the experiment compared to the expected result for a system without any spatial correlations.

In contrast, our results for co-culture assays suggest that great caution is warranted when calibrating mathematical models to replicate co-culture growth-to-confluence assays. We demonstrate that the spatial summary statistics for two species, applied in isolation, does not reliably indicate the suitability of a particular modelling framework when the two species are present in co-culture. Furthermore, we also show that extensions of the averaged agent coordination number data and spatial correlation index fail to provide a clear quantitative measure of the suitability of one model over another for a wide range of parameter space. In this regard, our results suggest that extreme care ought be exercised when interpreting co-culture experiments using mathematical models. Indeed, the results of this study suggest that further investigation of additional summary statistics that can reliably distinguish between the application of stochastic, mean-field and moment-dynamics modelling frameworks, is warranted.

In summary, our analysis indicates that it is possible to make a sensible distinction between the suitability of mean-field, moment-dynamics and stochastic descriptions of a growth-to-confluence assay for a mono-culture growth-to-confluence experiment by analyzing either the distribution of agent coordination number or the spatial correlation index. Conversely, for a co-culture growth-to-confluence experiment more care is needed, and neither of these summary statistics can reliably distinguish between the suitability of the three candidate modelling frameworks. To this end, we suggest that the most pragmatic approach to make an initial assessment of the parameters governing a co-culture experiment is to use the discrete model as a screening tool to obtain parameter estimates. Depending on these initial estimates, it may then be possible to implement either a mean-field or a moment-dynamics description of the system for further analysis, if required.

## Acknowledgements

We appreciate experimental support from Parvathi Haridas. This research is supported by the Australian Research Council (FT130100148) and the 2011 International Exchange Scheme funded by the Royal Society.

